# Considerations for clinical curation, classification and reporting of low-penetrance and low effect size variants associated with disease risk

**DOI:** 10.1101/556316

**Authors:** Ozlem Senol-Cosar, Ryan J. Schmidt, Emily Qian, Derick Hoskinson, Heather Mason-Suares, Birgit Funke, Matthew S. Lebo

**Author notes:** these authors contributed equally (Ozlem Senol-Cosar, Ryan J. Schmidt).

## Abstract

**Purpose:** Clinically relevant variants exhibit a wide range of penetrance. Medical practice has traditionally focused on highly penetrant variants with large effect sizes and, consequently, classification and clinical reporting frameworks are tailored to that variant type. At the other end of the penetrance spectrum, where variants are often referred to as “risk alleles”, traditional frameworks are no longer appropriate. This has led to inconsistency in how such variants are interpreted and classified. Here, we describe a conceptual framework to begin addressing this gap.

**Methods:** We used a set of risk alleles to define data elements that can characterize the validity of reported disease associations. We assigned weight to these data elements and established classification categories expressing confidence levels. This framework was then expanded to develop criteria for inclusion of risk alleles on clinical reports.

**Results:** Foundational data elements include cohort size, quality of phenotyping, statistical significance, and replication of results. Criteria for determining inclusion of risk alleles on clinical reports include presence of clinical management guidelines, effect size, severity of the associated phenotype, and effectiveness of intervention.

**Conclusions:** This framework represents an approach for classifying risk alleles and can serve as a foundation to catalyze community efforts for refinement.

## INTRODUCTION

Genetic variants that contribute to disease lie on a spectrum from rare alleles with large effect sizes to more common alleles with small effect sizes^1-3^. Genetic diseases have historically been categorized as either Mendelian (i.e. caused by variants at a single locus that segregate with a recognizable pattern within families) or complex (i.e. caused by a combination of several/multiple variants and environmental factors, often with some degree of heritability that does not follow a clear inheritance pattern)^4^. Accordingly, Mendelian variants are identified by studying affected families, whereas variants associated with common and complex disease are identified through association studies involving large populations of unrelated individuals^4^. This traditional, binary classification of disease was appropriate when the field focused on the most penetrant and severe heritable conditions but does not adequately describe the known landscape of heritable conditions that affect health today. It has long been known that many pathogenic variants do not always lead to disease when present in an individual (i.e. show reduced penetrance), but we are only now beginning to appreciate the enormous gradient of penetrance associated with variants that cause or contribute to genetic disease.

While the extreme ends of the penetrance and effect size spectrums are well described, clinically relevant variants in the “grey zone” between clear Mendelian and complex inheritance are ill-defined with regard to terminology, classification, and clinical reportability. These variants are found in the population more commonly than classic Mendelian alleles and can be inherited in sometimes recognizable familial patterns. Such variants are identified by both Mendelian case studies and population-based association studies and tend to be described using terminology depending on which type of study identified them. Mendelian frameworks refer to these variants as “low penetrance variants” and complex disease studies describe them as “risk variants” or “risk alleles”. For simplicity, we will use the term “risk allele” throughout this manuscript.

Some clinically significant risk alleles are well characterized and have long been included on clinical reports, but the lack of consensus terminology and interpretation criteria for this variant type has led to inconsistent classification^5^. Risk alleles can be observed at frequencies in population databases that meet Mendelian classification standards to be classified as “benign” ^6,7^. A well-known example is the *F5* p.Arg534Gln variant (Factor V Leiden), which is present in 3% of European alleles in gnomAD (https://gnomad.broadinstitute.org/variant/1-169519049-T-C, accessed 10/19/18) and has been submitted to ClinVar by 9 laboratories as “pathogenic” (n=4), “benign” (n=1) and “risk variant” (n=4) (ClinVar ID 642, accessed 10/19/18). Such divergent classifications can create confusion for patients and clinical practitioners.

With costs of genomic sequencing rapidly decreasing, large population-based studies are increasingly identifying such risk alleles^8,9^. Additionally, genomic screening is beginning to be offered to healthy individuals^10,11^ and there is increasing interest in returning these variants on clinical reports. As such, it is critical that the community defines frameworks for evaluating the validity of evidence supporting the role of risk alleles in disease and develops terminology to clearly distinguish these from variants that cause highly penetrant, Mendelian disease.

Furthermore, consensus is needed regarding what level of evidence warrants inclusion of risk alleles on clinical reports. While the scientific validity of the associated risk has to be the foundation, additional factors need to be considered to balance clinical utility and possible risks for unnecessary medical action. Some risk alleles have clear actionability, including those recognized by specific recommendations from professional societies. For example, the p.Ile1307Lys variant in the *APC* gene is associated with increased risk for developing colorectal cancer, especially in Ashkenazi Jewish population. The 2018 National Comprehensive Cancer Network (NCCN) guidelines recommend that unaffected individuals with this variant who are lacking family history of colorectal cancer begin colonoscopy screening at age 40, 10 years earlier than the general population (NCCN Genetic/Familial High-Risk Assessment: Colorectal Version 1.2018. https://www.nccn.org/professionals/physician_gls/pdf/genetics_screening.pdf. Accessed 01 October 2018). The NCCN guidelines caution providers to be aware of the changing landscape at this time and to consider patient preferences and new evidence that may present when implementing colonoscopy screening regimens. In contrast, reporting a variant that confers risk for a rare condition for which no effective preventative measures exist may require different consenting and counseling procedures, as the probability of developing disease and the severity of the impact may be difficult to convey and comprehend.

We compiled a set of representative risk alleles to define data elements that can be used to express the varying degrees of confidence in their ability to contribute to genetic disease. We propose a classification framework that is conceptually similar to the widely used American College of Medical Genetics and Genomics/Association of Molecular Pathology (ACMG/AMP) classification system for germline Mendelian disease^7^ and discuss which criteria may influence inclusion of such variants on clinical genetic reports. This work is intended to catalyze discussion in the genetics community and serve as a basis for refinement and standardization.

## MATERIAL AND METHODS

### Inclusion criteria

Variants included in this study were reported to be associated with clinically relevant phenotypes, as opposed to physical traits such as eye color and height. Additionally, these variants were supported by genetic association studies reporting statistically significant enrichment in cases versus controls.

### Variant set

Variants were selected from internal databases of two clinical genetic testing laboratories (Laboratory for Molecular Medicine and Veritas Genetics) as well as public databases such as ClinVar. Clinical tests performed by these laboratories include disease-focused gene panels, diagnostic exome/genome testing, and elective genomic screening. Selected variants for review ranged from single variants, multiple variants forming a haplotype, variants conferring risk through compound heterozygosity, and digenic risk variant combinations.

### Curation process and classification criteria

Data elements associated with these risk alleles were compiled and assigned weight depending on the strength of the evidence. This framework was applied to the variant set and refined to arrive at a final version (Figure 1). Variant classifications were performed by two independent curators. Evidence was gathered systematically using structured data collections forms (Supplementary Figure 1). Each classification was reviewed by ABMGG-certified clinical molecular geneticists and finalized after reaching consensus with the entire group.

**Figure 1.**
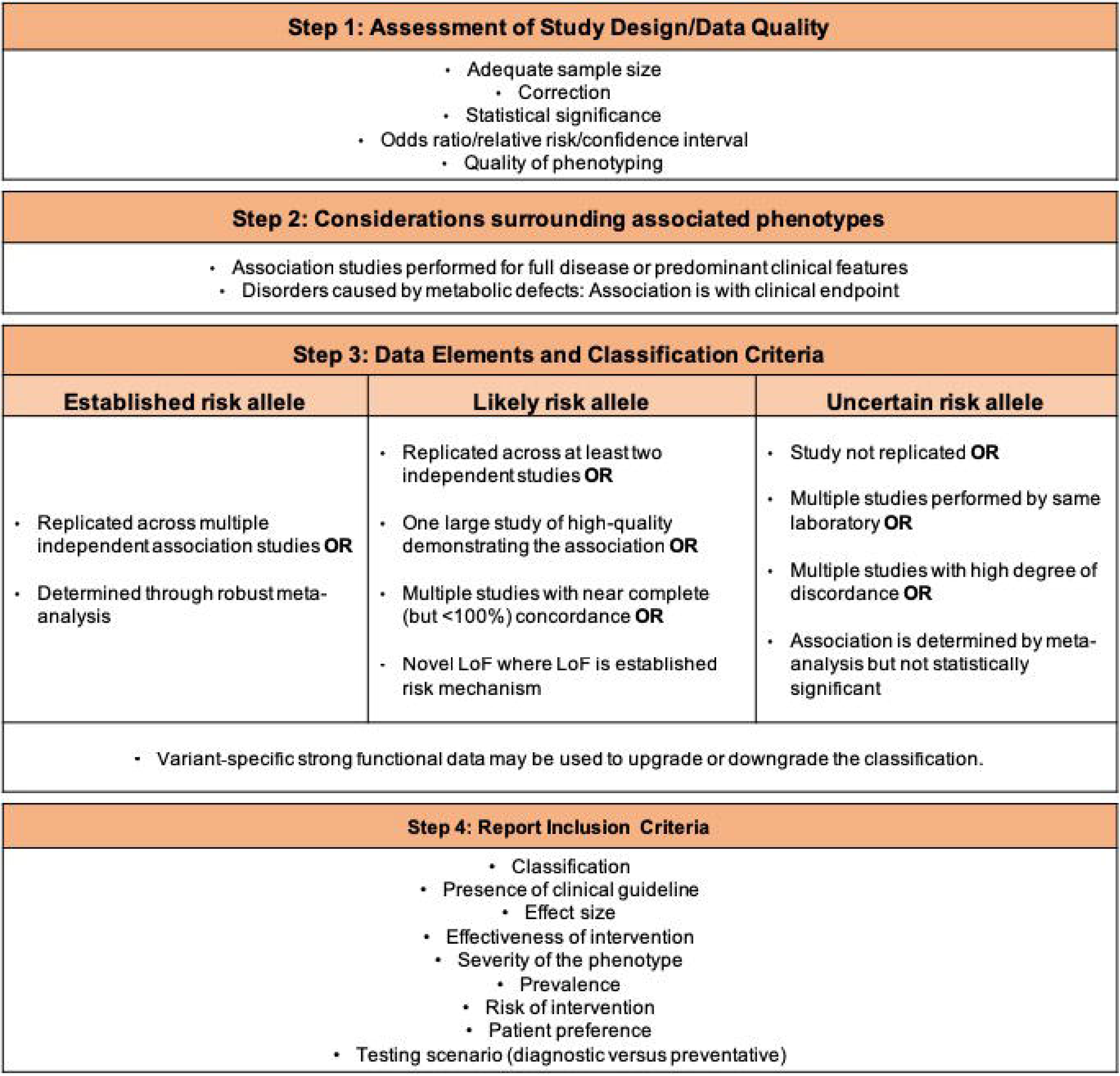
Decision-making framework for the classification of risk alleles

### Metrics for establishing reportability

We identified criteria for guiding decision making on whether or not to include risk alleles in clinical reports based on the level of the variant-disease association which was determined by the classification framework presented in this study, as well as general clinical information about the disease that is associated with the variant in question such as prevalence, severity, effectiveness of intervention, and risk of intervention. We extracted the guiding principles for these criteria from PubMed literature searches, Genetics Home Reference (https://ghr.nlm.nih.gov/), Center for Disease Control and Prevention (https://www.cdc.gov/), ACMG Technical Standards and Guidelines (http://www.acmg.net/ACMG/Medical-Genetics-Practice-Resources/Technical_Standards_and_Guidelines.aspx), Clinical Genome Resource Actionability Work Group documents (https://www.clinicalgenome.org/curation-activities/clinical-actionability/the-process/), disease-specific databases (e.g. https://www.nccn.org/professionals/physician_gls/recently_updated.aspx,) and other relevant publications^12-14^.

## RESULTS

### Classification framework

Classifying any type of variant in a clinical setting requires careful evaluation of the quality of the associated data, aggregation of available evidence, and application of criteria to establish the likelihood with which this evidence predicts the outcome. The following sections describe a proposed framework for assessing and classifying risk alleles using the terminology initially suggested by the ACMG/AMP guidelines for interpretation of germline sequence variants^7^. We intentionally focused on general steps and concepts (Figure 1 and sections below) to provide a basis for community iteration and refinement.

### Step 1: Assessment of study design and data quality

Characteristics of well-designed and reliable association studies have been published^15-17^ and include large, race-matched and well-phenotyped case and control cohorts, application of statistical correction for multiple hypothesis testing, application of a rigorous threshold for statistical significance, and calculation of odds ratios or relative risks as a measure of effect size. We considered any study that reported statistically significant results (p<0.05) and excluded those reporting effect sizes where the confidence interval included 1.

### Step 2: Considerations surrounding associated phenotypes

Medical literature often reports association of a variant across a range of phenotypes. While some represent distinct clinical entities, others represent endophenotypes, i.e. one of several features that together constitute a clinical entity and deciding which studies should be combined can be challenging for a non-expert. This is now a well-recognized phenomenon and early guidance is available to train clinical variant curation professionals (Guidance for lumping and splitting in ClinGen gene:disease clinical validity curations. https://www.clinicalgenome.org/site/assets/files/9703/lumping_and_splitting_guidelines_gene_curation_final.pdf. Accessed 10/01/18). While this problem affects variants across the full range of the genetic penetrance spectrum, it is particularly common among genetic association studies, which often examine a wide array of features ranging from the full disease to endophenotypes^18^. We intentionally took a very conservative approach to avoid over-classification of variants. Generally, only studies reporting an association with the full disease or its predominant clinical features were included. For example, for the *APOE* e4 allele we included studies that demonstrated an association with Alzheimer disease but not association with aggression or depression in Alzheimer patients nor with disease progression once an Alzheimer diagnosis was made.

Finally, there can be significant ambiguity as to what defines the disease state. This is particularly pronounced for disorders whose primary defect is a biochemical imbalance, which results in clinical features only when exceeding a threshold. For these disorders, we only considered studies reporting an association with clinically evident phenotypes. For example, hereditary hemochromatosis is caused by variants in the *HFE* gene that lead to elevated transferrin levels, which can eventually manifest with symptoms of end-stage organ damage secondary to iron storage. Because elevated serum transferrin alone can have different causes^19^, we considered only studies reporting an association with the clinical endpoint (e.g. liver disease).

### Step 3: Data elements and classification criteria

The strength of a variant-disease association was assigned one of three categories using terminology akin to those commonly used for Mendelian disease and suggested in the ACMG/AMP interpretation guidelines^7^: “established risk allele”, “likely risk allele” or “uncertain risk allele”. Because a large number of association studies fail to replicate^20^, highest emphasis was placed on meta-analyses and multiple, independent studies confirming the originally reported association. Additional data elements (such as functional data) were given modifying weight.

#### Number and types of studies reporting an association

To classify a variant as an “established risk allele” for a condition, we suggest a minimum of one robust meta-analysis or multiple independent case-control studies that each meet all criteria for well-designed and conducted studies outlined above. A “likely risk allele” classification requires less evidence and we suggest at least two independent case-control studies showing a statistically significant association with the phenotype of interest. When multiple studies report conflicting results, a “likely risk allele” classification can still be reached when the clear majority are concordant with regard to significance and effect size. Another scenario qualifying for a “likely risk allele” classification is a single, large study of high quality with data from multiple sites.

All other scenarios result in an “uncertain risk allele” classification as a baseline, which can be modified when other supporting or refuting evidence, such as functional data, is available (see below). Common examples for “uncertain risk allele” classifications include a single, unreplicated case-control study, replicated associations derived from overlapping cohorts, and replicated results derived solely from very small studies. Case-control studies that have already been included in meta-analyses are not individually reviewed and double counted as replication.

#### Functional data

Evidence demonstrating a direct effect on protein function was given supporting weight, allowing for adjustment of the classification category. Validity, relevance, and reproducibility of the functional data were taken into consideration as recommended by ACMG/AMP guidelines^7^. Only strong functional data was allowed to be used in this fashion^7,21^. Generally, this included only data from variant-specific *in vivo* models recapitulating the associated human phenotype or reliable enzymatic assays performed in relevant *in vitro* systems. Functional evidence of an effect on protein function can provide confidence in disease association and a distinct causal role for the variant, rather than an indirect effect through genetic linkage. In contrast, functional data was not used to downgrade the classification when association study results clearly support a “likely” or “established” risk allele classification as the signal can always be due to another variant that is in linkage disequilibrium. For example, even though *in vitro* functional studies demonstrated no effect on protein function, the p.Asn34Ser variant in *SPINK1* was classified as an “established risk allele” based on evidence from two meta-analyses and one large case-control study (Table 1).

**Table 1.**
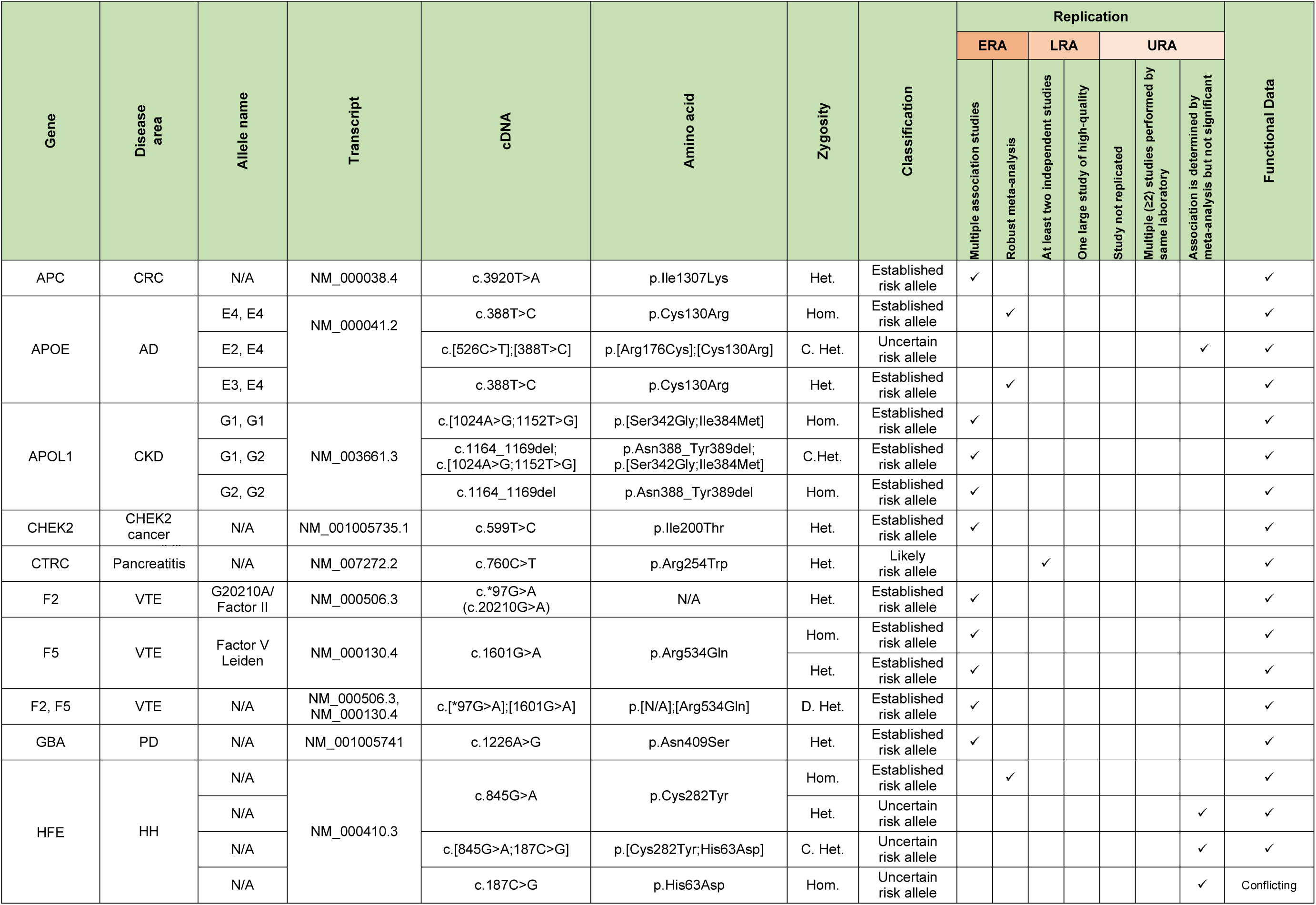
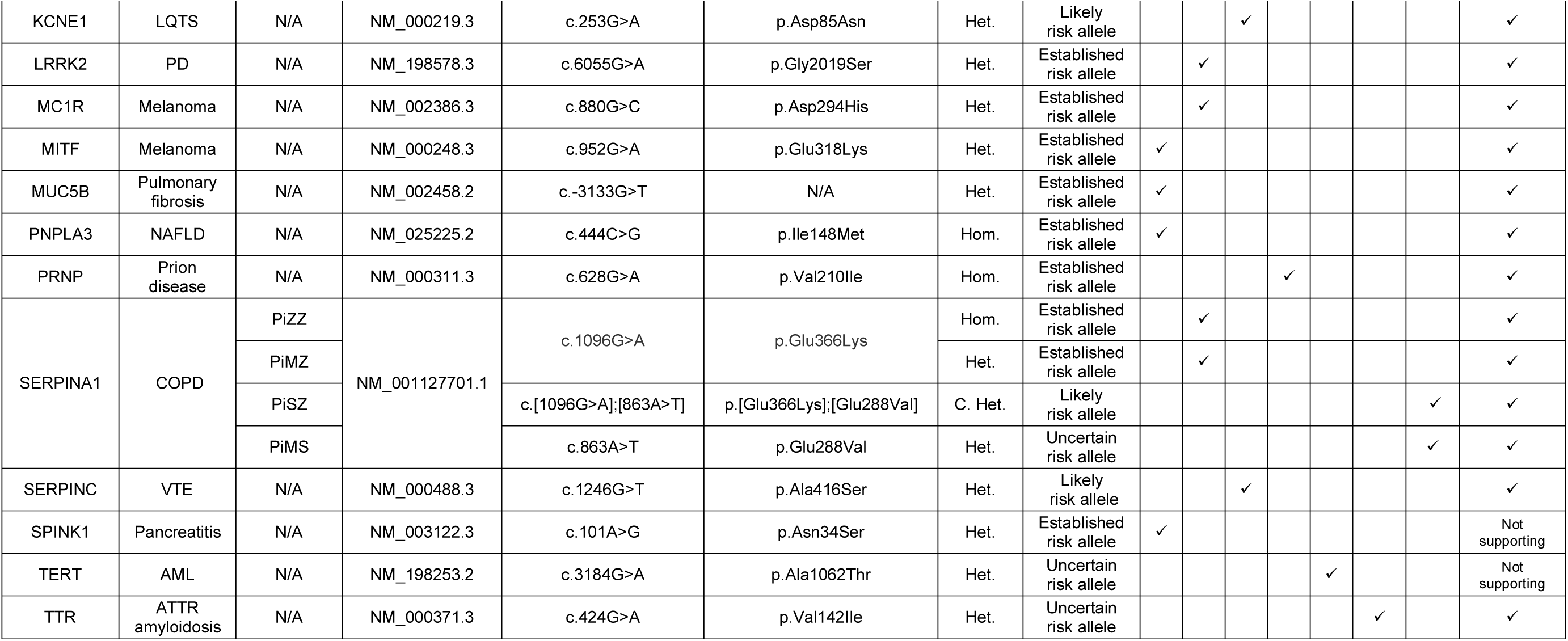
List of low-penetrant variants that were classified based on the framework presented in this study. CRC: colorectal cancer, AD: Alzheimer’s disease, CKD: chronic kidney disease, VTE: venous thromboembolism, PD: Parkinson’s disease, HH: hereditary hemochromatosis, LQTS: long Qt syndrome, NAFLD: nonalcoholic fatty liver disease, COPD: chronic obstructive pulmonary disease, AML: acute myeloid leukemia, ATTR: transthyretin, Het: Heterozygous, Hom: Homozygous, C. Het: Compound heterozygous, D. Het: Double heterozygous.

#### Loss of function variant considerations

Most risk alleles are common in the general population, which provides enough statistical power to establish an association with disease. However, many individual variants that confer risk may be rare, and thus would not have the power to be identified or established as risk factors. In some instances, it may be possible to risk calculations on a class of variants via aggregate variant association studies, something that is particularly possible for “loss of function” (LOF) variants. The ACMG/AMP Mendelian classification framework recognizes that when LOF is an established mechanism of disease for a gene, even novel LOF variants without any additional supporting evidence are attributed substantial weight. This concept can be extended to genes that are overall associated with lower penetrance. An example is the *CHEK2* gene, where LOF variants in general have been associated with an increased risk of cancer^22^. The most prominent cancer susceptibility variant in this gene is a LOF variant (c.1100delC) that leads to a 37% lifetime risk of cancer, which reaches a level where use of the Mendelian “pathogenic” classification under an autosomal dominant cancer susceptibility framework may be more appropriate^23^. However, because the *CHEK2* gene is not as well studied as other cancer susceptibility genes (such as *BRCA1* and *BRCA2*), it may be more prudent to describe novel LOF variants using the risk framework and elevate them to a Mendelian classification when more data is available as that conveys more certainty about the clinical outcome.

### Application of the framework

We applied this framework to a set of 33 variants in 22 genes that met characteristics to potentially be classified as risk alleles. Variants and alleles were assessed individually for disease associations across all zygosity states. Data was available to make classifications for 19 heterozygous variants, 9 homozygous variants, and 5 compound or double heterozygous variants. A summary of the criteria met and resulting classifications are listed in Table 1 with additional detail on each classification provided in Supplementary Table.

### 4. Reporting considerations

Deciding whether or not to return risk alleles in clinical genetic testing can be challenging as the absolute risk and the clinical utility of disease associations are often not as clear as they are for Mendelian disease variants. Here, we define criteria we believe should be taken into consideration for reporting decisions.

The base criterion for clinical reporting is the scientific validity of the associated risk, which is expressed by the classification of the variant. In our opinion, the clinical utility of returning variants with reported but unconfirmed disease associations (i.e. “uncertain risk alleles”) is low and we therefore propose to restrict reporting to “established” and “likely” risk alleles. This is similar to common practice for Mendelian testing, where predictive reports (secondary findings) are commonly restricted to “likely pathogenic” and “pathogenic” variants^9,24^. However, while a likely or established risk allele classification constitutes a necessary criterion, it is not sufficient. Below and in Figure 2 we describe additional criteria that should be considered when making decisions on including risk alleles on clinical reports.

**Figure 2.**
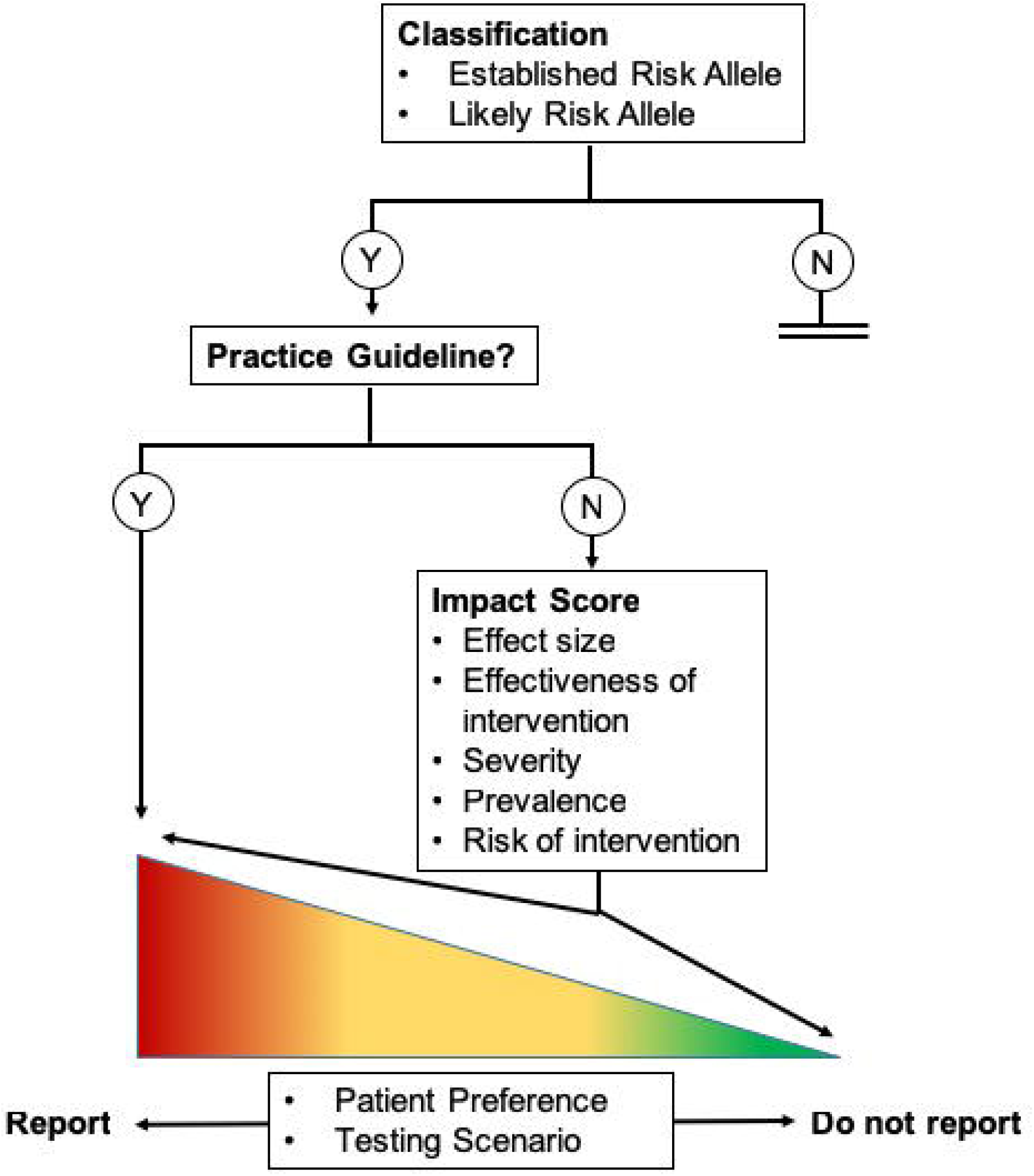
Routing logic for inclusion of risk alleles on clinical reports

A major consideration for returning a risk allele is the availability of clinical management or practice guidelines issued by expert groups or professional societies. Variants classified as established or likely risk allele with such guidelines were considered candidates to include on clinical reports in our framework. For other risk alleles, we discuss five additional clinical criteria that could be combined into an “impact score” reflecting the overall clinical importance of the finding: a) effect size, b) disease prevalence, c) disease severity, d) effectiveness of intervention, and e) risk associated with action/intervention. The scores for effectiveness of intervention, severity, and risk of intervention were based upon the semi-quantitative metrics put forth by the ClinGen Actionability Working Group^13^. As genomic testing is increasingly administered in an elective fashion (often referred to as consumer genomics), personal utility and testing scenario (diagnostic versus predictive) should also be considered.

To arrive at such an impact score we assigned a numerical value from 0 to 3 to each criterion (3 having the greatest weight) and applied this system to three illustrative risk alleles. Table 2 lists the rubric for assigning the scores and Figure 3 shows the scores obtained for three representative risk alleles: p.Asp85Asn (*KCNE1*), p.Glu318Lys (*MITF*) and p.Val210Ile (*PRNP*) variants. Please note that each criterion that we proposed had equal weight on the impact score. The total scores were calculated for comparison purposes between the variants with variable clinical impact. Even though we propose criteria to calculate clinical impact, further consideration by the broader community would be required to determine a universal threshold for reportability.

**Table 2.** Scoring system used to assess the strength of the criteria for reportability

|  | **0** | **1** | **2** | **3** |
| --- | --- | --- | --- | --- |
| **Effect size** | OR=1 | OR 1-2 | OR 2-4 | OR>4 |
| **Effectiveness of Intervention** | Controversial or unknown effectiveness | Minimally effective | Moderately effective | Highly effective |
| **Severity** | Minimal or no morbidity | Modest morbidity | Possibility of death or major morbidity | Reasonable possibility of sudden death |
| **Prevalence** | Unknown | Rare (1:2000) | Intermediate (1:100-1:2000) | Common >1:100 |
| **Risk of intervention** | Highly risk, poorly acceptable or intensive interventions **OR** no known intervention | Moderate risk, moderately acceptable or intensive interventions | Greater risk, less acceptable and substantial interventions | Low risk, or medically acceptable and low-intensity interventions |

**Figure 3.**
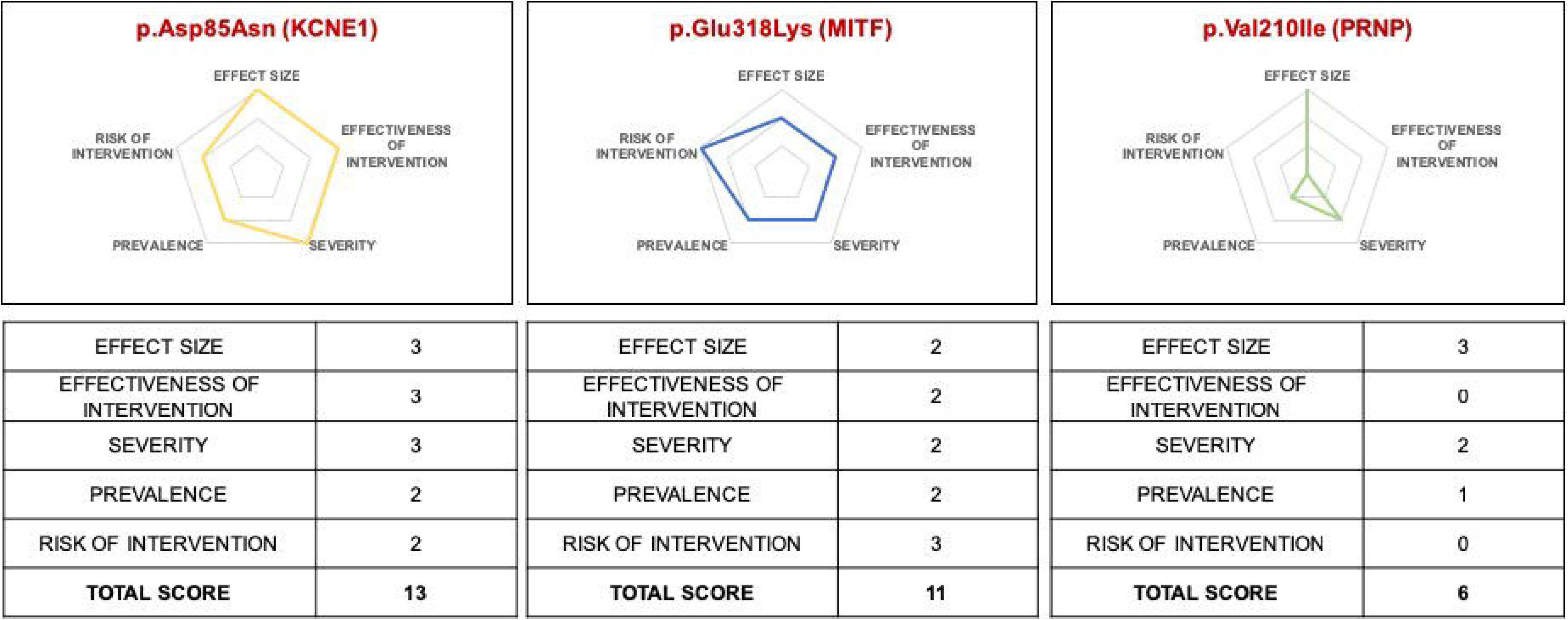
Reportability scores. Radar charts visualize 5 reportability criteria for three variants

### Considerations for communicating the significance of risk on a clinical report

Similar to what is customary for reporting Mendelian disease variants, risk alleles should be accompanied by a summary of all evidence supporting a classification. If an association study reports a statistically significant odds ratio (or other statistical measure), these values along with confidence intervals and p-values should be stated or summarized. Additionally, it is important to point out when findings were limited to populations of a specific ancestry. In addition, as statistical measures derived from this framework do not represent absolute risks, care needs to be taken to communicate this clearly and avoid over-interpretation by the recipient. A sample interpretive summary is provided below.

“F5 c.1601G>A (p.Arg534Gln; commonly known as Factor V Leiden, historically reported as p.Arg506Gln) has been associated with increased risk for venous thromboembolism (VTE). This variant has been observed in multiple ethnic backgrounds with highest frequencies in individuals of European ancestry (2.96%, Genome Aggregation Database (gnomAD); rs6025) and is present in ClinVar (ID: 642). Several meta-analyses and case-control studies have reported odds ratios between 2.2-4.93 for developing VTE in heterozygous carriers (OR=2.2 [95% CI 2.0-2.5]^25^; OR=4.22 [95% CI 3.35-5.32]^26^; OR=4.93 [95% CI 4.41-5.52]^27^; OR= 2.4 [95% CI 1.3– 3.8]^28^) and odds ratios between 7-11.5 for developing VTE in homozygous carriers (OR=7.0 [95% CI 4.8-10] Sode 2013; OR=11.45 [95% CI 6.79-19.29] Simone 2013). In vivo and in vitro functional studies provide evidence that the Factor V Leiden variant impacts protein function^29-32^. In summary, the p.Arg534Gln variant meets criteria for classification as an established risk allele for VTE.”

## DISCUSSION

The medical community has long been aware of variants that fall into the grey zone between rare, highly penetrant variants and variants contributing to common or complex disease. While the extreme ends of this penetrance gradient are clinically well defined, little guidance exists for variants that have significantly reduced penetrance yet have effect sizes that warrant consideration in a clinical setting. As was the case for Mendelian variants, the lack of standards has led to discordance in how these variants are evaluated and labeled, which may ultimately have negative consequences if they are classified as “benign” and not reported to the patient. As genomic testing is shifting towards exome and genome sequencing, the availability of large datasets is increasingly revealing risk alleles that are associated with medically relevant conditions. Simultaneously, the rise of elective genome screening in healthy individuals is increasing the demand to return such variants on clinical reports.

To address the emerging need for guidance, we developed a proposed first framework to systematically evaluate the scientific validity of reported risk allele associations. We define the data elements that should be evaluated and suggest a weighted method to assign clinical classifications. The utility of such frameworks is well established and is known to lead to harmonization between clinical laboratories. This is most recently evidenced by the enormous impact of the ACMG/AMP variant classification framework for Mendelian variants^7^, which has become widely used since it was first published in 2015 and which has led to an impressive amount of community harmonization aided by the availability of the ClinVar database and community efforts such as the Clinical Genome Resource (ClinGen)^5,33,34^. Additionally, we raise the question as to which additional factors should guide the inclusion of such variants on clinical reports. Scientific validity is the minimum requirement, but even more than for Mendelian disorders, the risk for over-interpretation by the recipient and the resultant risk for causing harm and anxiety has to be carefully considered for risk alleles.

Our approach was deliberately conservative and was designed to raise concepts rather than suggest prescriptive guidance. The latter will require iteration via community input and ultimately professional society recommendations. Similar to the ACMG/AMP classification framework, there is a need for developing disease and gene specific derivatives as expert knowledge is critical to better define the weight that can be assigned to certain data elements and to guide decision making for including credible risk alleles on clinical reports. Whether or not to return risk alleles will also be impacted by the testing scenario. We predict that the “bar” for including risk alleles may be lower in a diagnostic setting compared to a healthy/elective testing scenario where one may consider including uncertain risk alleles relevant to the indication, similar to what is common practice in traditional, Mendelian space^9^.

Finally, the framework we present here can be extended to protective alleles, which have been largely ignored in a traditional clinical testing setting but are expected to increase in demand as genomic testing is further expanded outside of the diagnostic context into the prediction of disease risk. Future work will be required to develop consensus approaches for accurate calculation and communication of disease risk based on the presence of risk and protective alleles. One specific challenge is to determine from which particular study the associated effect size, or magnitude of risk, should be derived. In general, larger well-controlled studies provide an effect size measurement that is closer to the true value for the entire population. Also, consensus approaches should be developed to accurately communicate the uncertainty surrounding any quantitative risk estimate. It will be important to avoid large discrepancies in risk estimates between clinical laboratories, which have previously plagued the direct-to-consumer testing space^35^.

As stated in the results section, the penetrance threshold that separates variants that should be classified by the Mendelian framework from risk alleles has not been firmly established. Consistently applying the appropriate framework may be challenging given the lack of penetrance estimates for most variants. Additionally, different disease areas within clinical genetics may wish to apply different penetrance thresholds. For example, cancer predisposition testing, which includes the *CHEK2* c.1100delC variant, has a longstanding history of classifying variants within the Mendelian framework despite incomplete penetrance for many variant-cancer type associations. In our opinion, as a general rule when penetrance data is unavailable, variants whose disease association has been demonstrated through only segregation analysis within affected families should be classified within the Mendelian framework whereas variants identified in association studies or case-control cohorts of unrelated individuals should be classified as risk alleles.

This work focuses on the classification and reporting of risk alleles whose association with disease is clinically significant in isolation, ignoring the potential effects of additional genetic and environmental variables that impact disease risk. Our classification framework can be applied to any individual variant or genotype regardless of the magnitude of risk that they are associated with. Laboratories may choose different thresholds for reporting risk alleles that will impact the number of these variants that appear on clinical reports.

Given the potential predictive utility inherent in genetic testing, risk (and protective) alleles will play an important role in the generation of clinically relevant quantitative risk estimates as the full spectrum of genetic and environmental variables that influence disease risk becomes more well understood. The classification criteria presented here may serve as a starting basis for inclusion in models of disease risk such that only those variants classified as established or likely risk alleles are included. Future challenges in developing clinically relevant quantitative risk estimates include: 1) calculating the patient’s pre-test risk based on demographic, clinical, environmental, and other genetic variables, 2) selecting and integrating the published data used to calculate the updated risk estimate, 3) updating the risk model in an accurate and transparent manner, 4) appropriately expressing uncertainty surrounding the calculated risk estimate in the clinical report.

Our work serves as a starting point for the structured classification and reporting of risk alleles in clinical molecular diagnostic reports. We look forward to continued advancement and harmonization on this subject within the broader clinical genetics community in the years to come.

