## Supplementary Figure 1 for "Considerations for clinical curation, classification and reporting of low-penetrance and low effect size variants associated with disease risk"

**Supplementary Figure 1**: Representative demonstration of a data-collection form used for literature curation.

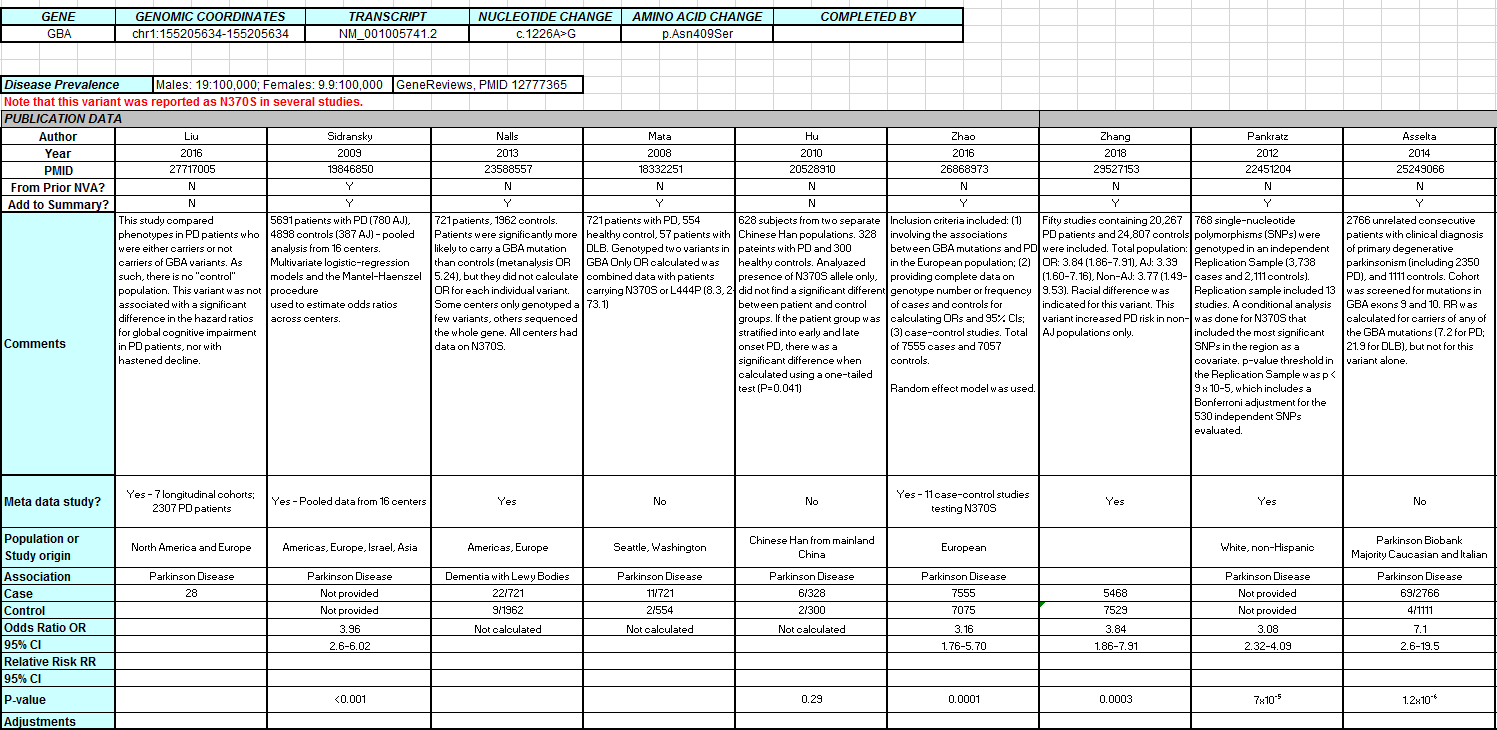
